# Wide dynamic range of contrast-encoding in neural responses in macaque V1

**DOI:** 10.64898/2026.06.15.732419

**Authors:** Hiroichi Yoshida, Yuzhi Chen, Wilson S Geisler, Eyal Seidemann

**Author notes:** Corresponding author email addresses: Hiroichi Yoshida, Eyal Seidemann.

## Abstract

The dynamic range of contrast-encoding in the early visual system has been investigated at both single-neuron and population levels in animals using oriented stimuli such as gratings and Gabor patches. However, contrast-encoding of unoriented stimuli such as Gaussians has been less explored, even though such stimuli can evoke a large population response. In studies that employ Gaussians, contrast response functions (CRFs) of neurons and neural populations in primary visual cortex are typically characterized across a limited range of Weber contrasts up to 100%, which may not adequately reflect the statistics of contrast in natural scenes. Indeed, our analysis shows that locations with contrasts far exceeding 100% Weber contrast are ubiquitous in natural scenes. Thus, our current understanding of contrast encoding remains incomplete in the context of natural environments. Here, we measured spiking activities of individual neurons using electrophysiology and population responses using calcium imaging in V1 of fixating macaques while Gaussian stimuli were presented over a wide range of Weber contrasts, up to 900%. Both the average spiking response and the average population response continued to increase robustly above 100% Weber contrast. Only a small minority of neurons saturate below 100% Weber contrast. These results demonstrate that the dynamic range of contrast-encoding in V1 is broader than previously assumed and aligns more closely with the statistics of contrast in natural scenes.

**Significance Statement:** The dynamic range of contrast-encoding in V1 has typically been characterized using only a limited range of contrasts, often excluding the high contrasts commonly found in natural scenes. We measured contrast response functions of individual neurons and neural populations in V1 across a much wider range of contrasts. We found that the responses of the majority of neurons, as well as the overall population, increased robustly up to extremely high contrasts. These findings suggest that the dynamic range of contrast encoding in V1 extends well beyond the commonly tested range.

## Introduction

A fundamental goal of visual neuroscience is to understand how key properties of natural scenes are encoded by the visual system. One such property is contrast. Since the spatial pattern of contrast remains relatively invariant to variations in uniform illumination, it is used by the brain to encode and recognize specific patterns of surface reflectance and shadow across different lighting conditions. For instance, responses of retinal ganglion cells (RGCs) to contrasts are largely invariant to background luminance (Sakmann & Creutzfeldt, 1969). Transforming photoreceptor responses into contrast responses early in the visual system reduces the range of values that need to be encoded in subsequent processing. Contrast signals contain most of the useful information in natural scenes but generally have a much smaller range than luminance across a scene and time of day.

The dynamic range of contrast-encoding by neurons is typically assessed by measuring a contrast response function (CRF). Because most V1 neurons are orientation-selective, the CRFs of single-cells and neural populations in V1 have been studied primarily using Gabor and grating stimuli (Albrecht and Geisler, 1991; Albrecht & Hamilton, 1982; Albrecht et al., 2002; Geisler and Albrecht, 1997; Palmer et al., 2017; Seidemann et al., 2016; Sit et al., 2009). However, V1 neurons also respond to unoriented stimuli, such as Gaussians. While Gabors strongly activate a select subpopulation at a given retinotopic location, Gaussians moderately activate almost all of the neurons at that location. Indeed, imaging results in macaque V1 suggest that Gabors and Gaussians of equivalent size and maximal luminance produce similar magnitudes of V1 population response, and are equally detectable by macaques (Chen et al., 2012), consistent with findings in humans (Watson and Ahumada, 2005). Furthermore, natural stimuli are composed of mixtures of narrow– and broad-band stimuli such as Gabors and Gaussians, respectively. Therefore, it is important to characterize the CRFs of V1 neurons to Gaussians in addition to Gabors.

Most behavioral and physiological vision experiments add visual stimuli to a uniform grey background having half the maximum display luminance. This prevents the maximum luminance of an added stimulus from exceeding twice the background luminance, and thus limits contrast to less than 100% Weber contrast, defined as:

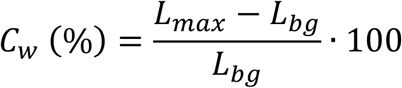

CRFs of V1 neurons or neural populations measured in this way often start to saturate or reach saturation at Weber contrasts lower than 100% (Chen et al., 2022; Kumar et al., 2019). However, the statistics of local Weber contrasts in natural scenes have not been studied systematically, leaving open the possibility that bright patches with luminance that is more than twice that of their background are common in natural scenes. If that were the case, then V1 neurons and neural populations might encode contrasts well beyond the previously explored range.

We quantified local contrasts in natural images and found that patches exceeding 100% Weber contrast are indeed common. This observation raises the plausible hypothesis that V1 neural populations continue to increase their response to stimuli with Weber contrast above 100%, which is beyond the previously explored range. Consistent with this hypothesis, our recent study showed that a direct low-power optogenetic stimulation in V1 can evoke a significantly higher population response than a Gaussian of 100% Weber contrast (Chen et al., 2022). Together, these findings highlight the importance of measuring V1 responses to a wider range of contrasts.

To measure V1 responses to a wide range of Weber contrasts, we devised a method to generate Gaussian stimuli with Weber contrasts up to 900%. We measured single– and multi-unit spiking activities, and widefield fluorescent calcium imaging signals (GCaMP) in macaque V1 during a fixation task. We show that both the average unit response and the average GCaMP response continue to increase robustly above 100% Weber contrast, and that only a small minority of V1 neurons saturate below 100%. Consequently, the dynamic range of V1 population responses to localized stimuli is substantially wider than previously assumed and more consistent with the statistics of contrast in natural scenes.

## Results

### Initial exploration of the range of local contrasts in natural scenes

To begin exploring the range of local contrasts in natural scenes, we computed what we call the “center-surround contrast” (*C_cs_*) at every pixel location in a set of five natural scene images by convolving them with kernels consisting of concentric center and surround subregions (**Figure 1A**). In each image, *C_cs_* at a given image pixel location (*x*, *y*) was computed by placing the kernel such that its central pixel aligned with the image pixel, and taking the mean luminance of the image pixels falling within the center (*L_c_*_(*x,y*)_) and the surround (*L_s_*_(*x,y*)_) regions as follows:

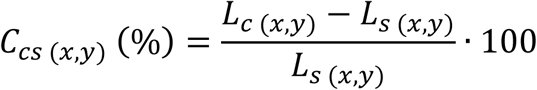

**Figure 1.**
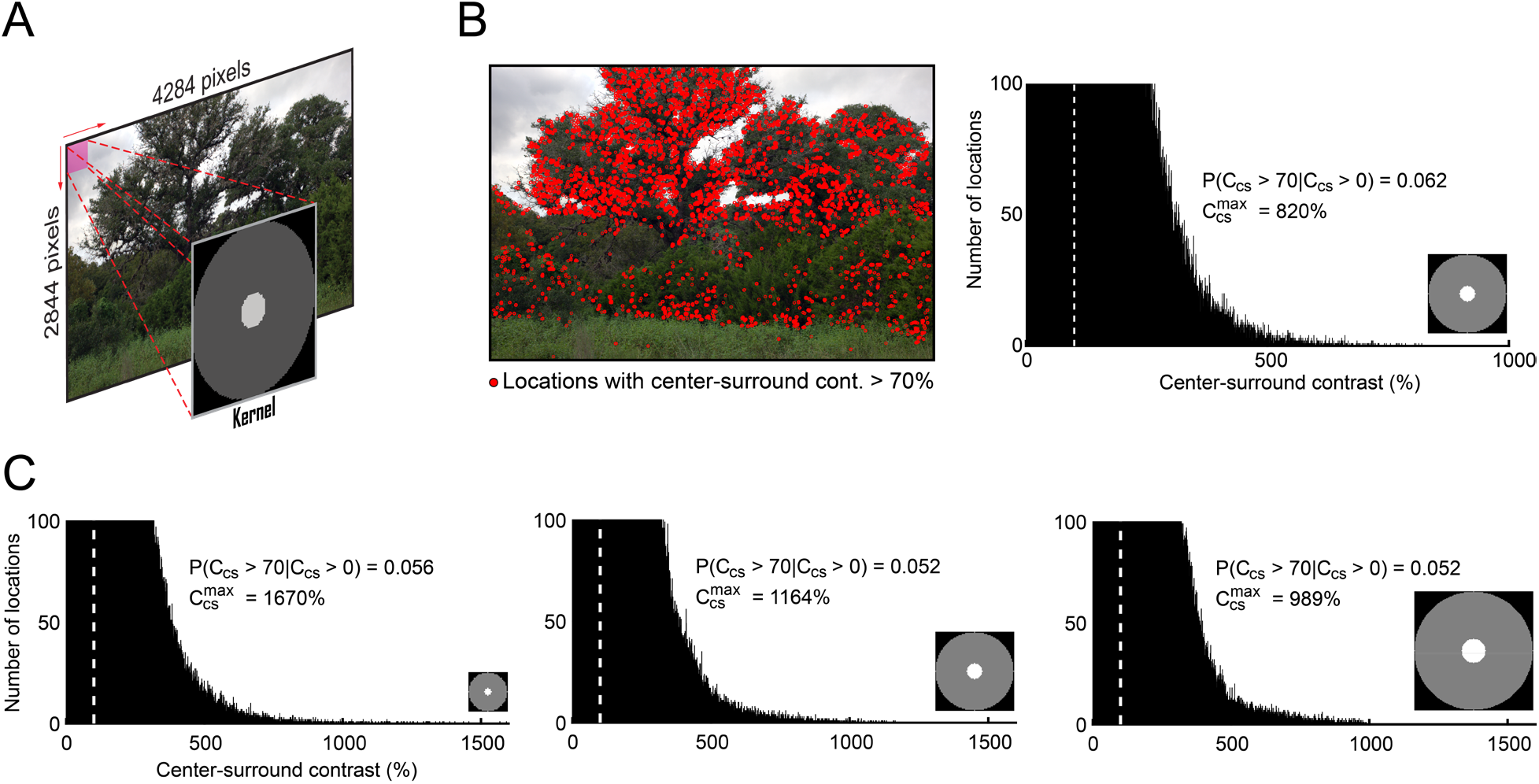
Distribution of center-surround contrasts in natural scenes. *A,* Contrasts in natural scene images (50 px/deg) are computed by convolving them with a kernel that has concentric center and surround regions (see Methods). ***B,*** *Left*, an example natural scene image. Red circles mark the spots with a center-surround contrast greater than 70%. *Right,* distribution of the center-surround contrasts in the image. Inset shows the kernel used to convolve the image (15:71 px for the center and the surround diameters, respectively). ***C,*** Distribution of the contrasts pooled across all images (N = 5), computed by kernels of different sizes (7:35, 15:71, and 29:141 px for the center and the surround diameters, respectively, from left to right).

The surround diameter was set to be five times larger than the center diameter to roughly match the large surround receptive field (RF) size of RGCs (Croner and Kaplan, 1995). We chose this definition because it is closer to the contrast signal encoded by neurons in the retina and the lateral geniculate nucleus (LGN) (Enroth-Cugell and Robson, 1966; Hubel and Wiesel, 1961; Kuffler, 1953).

A Gaussian stimulus of 100% Weber contrast has the maximal center-surround contrast of 70% when convolved with a kernel having a center radius equal to one standard deviation of the Gaussian stimulus (i.e., larger and smaller kernels give a lower center-surround contrasts). **Figure 1B** shows the locations in an example natural scene image where the center-surround contrast exceeds 70%. Such locations are ubiquitous (approximately 5% of the entire distribution), with the maximum center-surround contrast reaching as high as 821%. These results are robust to different images and kernel sizes, demonstrating that patches with high contrast are common in natural scenes (**Figure 1C**). Building on this result, our next goal was to determine how individual neurons, and particularly neural populations, in V1 respond to stimuli across such a wide range of contrasts.

### Contrast responses of single– and multi-units in V1 show a wide range of nonlinearities

Our main goal was to measure the neural responses in V1 to a range of Weber contrasts that exceed 100% and to understand vision under more natural conditions. To create Gaussians with such extremely high contrasts, we reduced the mean background luminance of the CRT monitor to 10 cd/m^2^, which was 10% of its maximal luminance. This approach allowed the maximal luminance of the stimulus placed on the background to reach up to ten times larger than the background luminance, achieving the maximal Weber contrast of 900% (**Figure 2A**), while keeping the mean luminance in the photopic range. We recorded the activity of individual neurons, the constituents of the population code, and the activity of neural populations in V1 while two animals performed the fixation task where a small white Gaussian (0.33° FWHM) was flashed briefly over two temporal cycles at the corresponding retinotopic coordinates of the recording site (see Methods; **Figure 2B**). The stimulus contrast varied pseudo-randomly from trial to trial between 0-900%.

**Figure 2.**
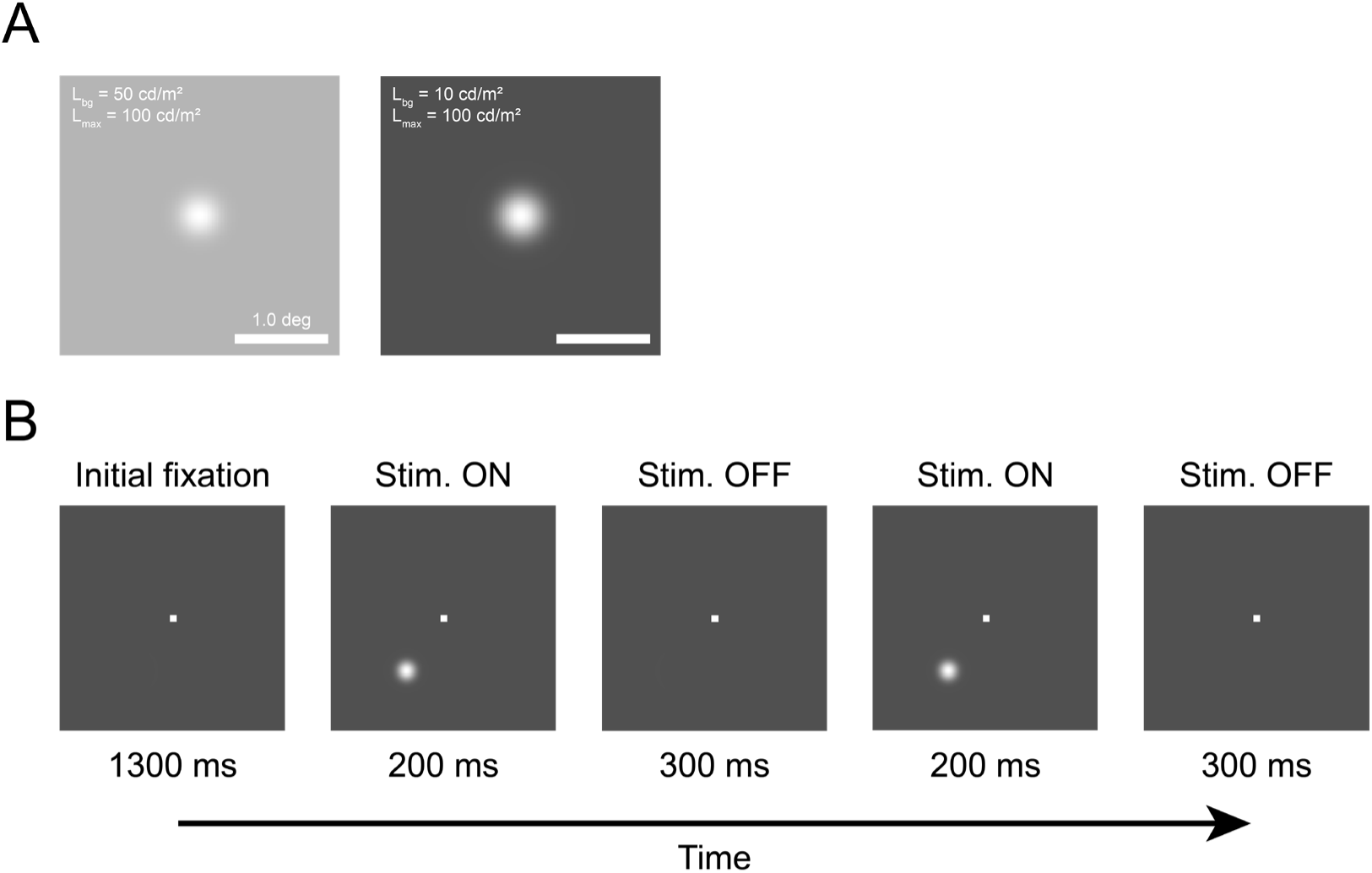
Stimulus and behavioral paradigm. *A, Left,* a small white Gaussian stimulus (0.33° FWHM; 50 px/deg) of 100% Weber contrast placed on a uniform grey background (50 cd/m^2^). *Right,* a Gaussian stimulus of 900% Weber contrast placed on a uniform dark grey background (10 cd/m^2^). ***B,*** Fixation task. Following the initial fixation period, a Gaussian stimulus is presented at the receptive field (RF) of the electrode’s penetration site or the imaging site for two cycles. The monkeys maintain fixation at the central dot over two cycles of stimulus presentation to receive a reward.

To characterize the dynamic range of individual neurons, we recorded single– and multi-unit spiking activities using multi-channel probes (a 32-channel Plexon probe and a 384-channel NHP Neuropixels probe). The data were spike-sorted offline and the CRFs of all the identified units were fitted independently with the Naka-Rushton function that has two key free parameters, the power exponent, *n_s_* and the semi-saturation constant, *C*_50_ (see Methods). We imposed three criteria on the fits of the CRFs to select the units with valid visual-evoked responses to the stimulus. We only included the units that passed the criteria for the analysis below (**Table 1**). However, we later show that our key results hold even without any exclusion (**Figure 7A**). To quantify the contrast where the CRFs approach response saturation we computed, from the fitted Naka-Rushton curve, the contrast, *C*_90_ where response reached 90% of maximum, *C_max_*.

**Table 1.**
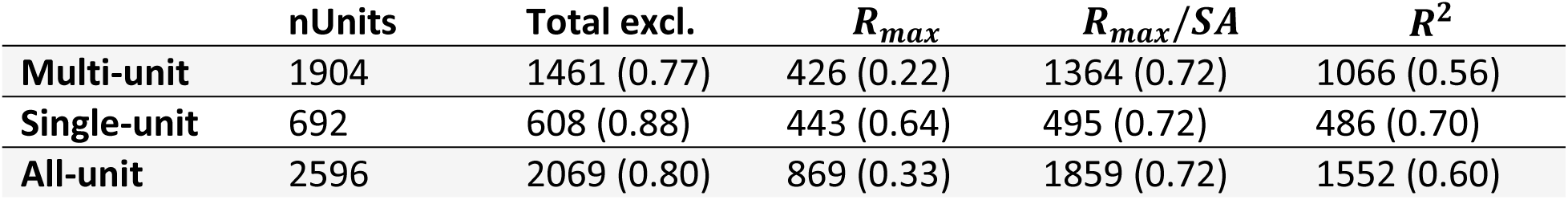
Number and proportion of excluded single– and multi-units by criterion. Three criteria were imposed on the fitted CRFs of all the single– and multi-units identified by spike-sorting, based on the fitted maximum firing rate, *R_max_* and spontaneous firing rate, *SA*, and the coefficient of determination, *R*^2^, to select units with reliable visual-evoked responses for analysis (see Methods). Numbers in the brackets indicate the proportion with respect to the multi-, single– or all-unit category.

| | nUnits | Total excl. | $R_{max}$ | $R_{max}/SA$ | $R^2$ |
| --- | --- | --- | --- | --- | --- |
| **Multi-unit** | 1904 | 1461 (0.77) | 426 (0.22) | 1364 (0.72) | 1066 (0.56) |
| **Single-unit** | 692 | 608 (0.88) | 443 (0.64) | 495 (0.72) | 486 (0.70) |
| **All-unit** | 2596 | 2069 (0.80) | 869 (0.33) | 1859 (0.72) | 1552 (0.60) |

Within our selected pool of single– and multi-units, a subset of the units had a high *n_s_* (> 2) and a low *C*_50_ (< 100). The spike response in these units started to saturate well below 100% contrast and *C*_90_ was smaller than 100%. An example of such a unit is shown in **Figure 3A and B**. However, in another subset of units with a low *n_s_* (≤ 1) and a high *C*_50_ (> 100), the response continued to increase robustly beyond 100% contrast, and in many of these cells, the response continued to increase all the way up to 900% contrast with their *C*_90_ projected to be greater than 900%. An example of such a unit is shown in **Figure 3E and F**. For the units with a low *n_s_* (< 1) and a low *C*_50_ (< 100), the response did not start to saturate by 100% contrast but did not necessarily increase robustly beyond 100% contrast either, with their *C*_90_ estimated to be greater than 100%. An example of such a unit is shown in **Figure 3C and D**. These observations suggest that single– and multi-units in V1 exhibit a wide range of nonlinearities in their contrast responses and that some V1 neurons respond robustly to contrast above 100%.

**Figure 3.**
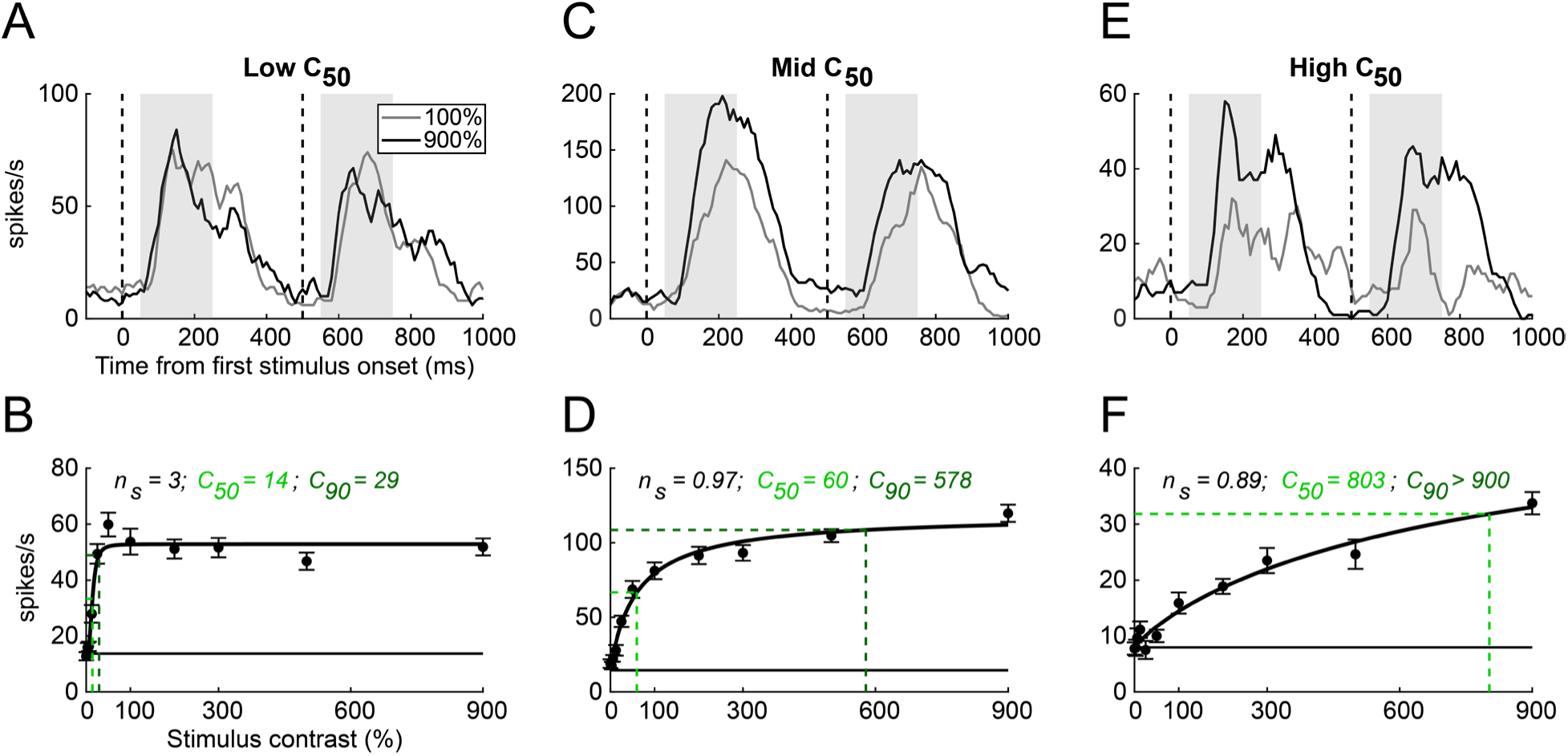
Example multi-unit responses in V1 across a wide range of visual contrasts. *A,* Trial-averaged peristimulus time histogram (PSTH) of an example multi-unit with a low *C*_50_ for 100% and 900% contrast stimuli, aligned to the first stimulus onset. Spike count in the preceding 50 ms window is scaled to spike rate (number of spikes per second) for every 10 ms for visualization. Vertical dashed lines indicate the stimulus onsets. Shaded regions indicate the time windows (50–250 ms after each stimulus onset) over which the spike rate is computed for each trial to generate the CRF shown in ***B***. ***B,*** Trial-averaged CRF of the multi-unit shown in ***A***, fitted by a Naka-Rushton curve. Error bars indicate SE across trials (n = 20). Dashed light and dark green lines indicate the responses at *C*_50_ and *C*_90_, respectively. Horizontal solid line indicates the fitted spontaneous firing rate, *SA*. ***C-D*,** Same as ***A-B*** for a multi-unit with a mid *C*_50_. ***E-F*,** Same as ***A-B*** for a multi-unit with a high *C*_50_.

### Majority of V1 units’ responses rise robustly above 100% Weber contrast

Given the variability in the response saturation profiles of our sample of V1 single– and multi-units, we next evaluated the distribution of the three parameters summarizing the shape of the CRFs, *n_s_*, *C*_50_, and *C*_90_ across the population. The two fitted parameters, *n* and *C*_50_ were revealed to share a systematic relationship that can be described by an exponential decay function, and their distributions were independent of the unit type (**Figure 4A**). The majority of the selected units were revealed to have *n* smaller than 2. Most importantly, cells that approach their response saturation below 100% Weber contrast (i.e., cells with *C*_90_ < 100) occupied less than 10% of our sample of single– and multi-units (**Figure 4B**). The majority of the units had *C*_50_ and *C*_90_ greater than 100%, producing CRFs similar to the example units with a low *n*_*c*_ (< 1) or a high *C*_50_ (> 100) (**Figure 3D and F**).

**Figure 4.**
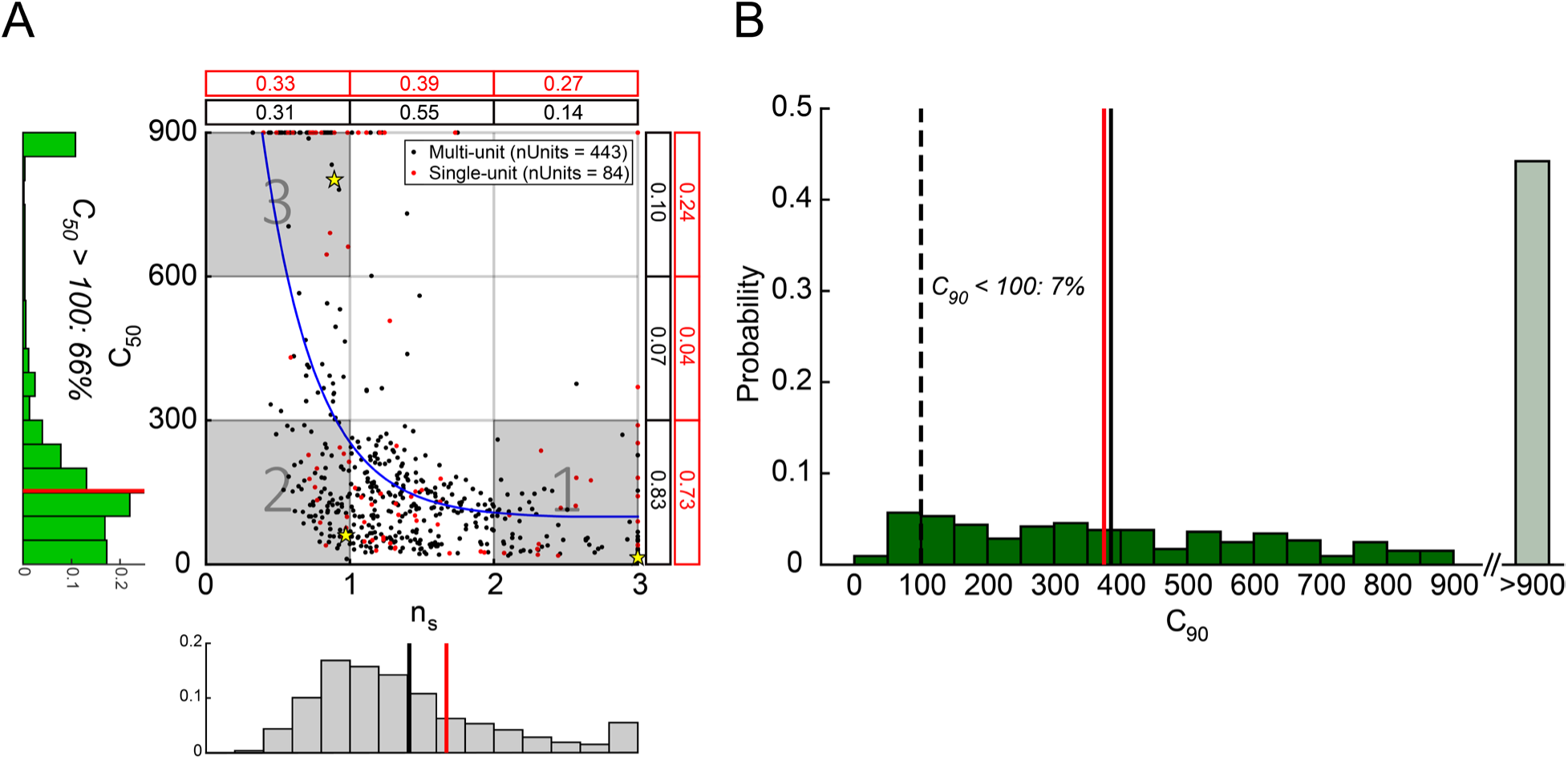
Summary of single– and multi-unit data. *A*, *C*_50_ as a function of *n_s_* for the selected single– (red) and multi-units (black) is fitted by an exponential decay function (blue curve; *k*_1_ = 2290.65, *k*_2_ = 97.16, *m* = 2.68; see Methods). Vertical solid lines on the marginal histograms indicate mean values for single– (red; 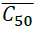= 153, 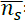 = 1.67) and multi-units (black; 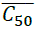 = 151, 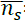 1.41) after excluding those with *C*_50_ > 900. Decimals on the top and the right edges of the figure indicate the proportion of units in each trisected subpopulation of *n_s_* and *C*_50_ for single– (red) and multi-units (black). Yellow stars indicate the three example units shown in Figure 3, each drawn from one of the three bins (grey shaded regions). ***B*,** Distribution of *C*_90_ derived from the CRFs fitted to the selected single– and multi-units. Vertical solid lines indicate mean values for single– (red; 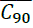 = 375) and multi-units (black; 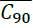 = 386) after excluding those with *C*_90_ > 900.

We binned these units into three groups (grey shaded regions in **Figure 4A**) based on their *n_s_* and *C*_50_, each of which includes one of the three example units shown in **Figure 3**. Consistent with the example multi-units, the within-bin average response reached a much higher peak at 900% contrast compared to 100% contrast across the two stimulus cycles for Bin 2 and 3, but not in Bin 1 (**Figure 5A, D and G**). To examine the dynamics of the response across the bins, we focus on the normalized within-bin average responses in the first cycle. The latency of the response increased, and the steepness of the rising phase decreased with increasing bin number (**Figure 5B, E and H**). The latency was shorter at 900% contrast compared to 100% contrast in all the three bins. While the average CRF within Bin 1 had only its *C*_90_ exceed 100% contrast, the average CRFs within Bins 2 and 3 had both of their *C*_50_ and *C*_90_ above 100% (**Figure 5C, F and I**). For Bin 3, with its *C*_50_ fitted to be larger than 900%, the average response continued to increase through 900% contrast without an immediate sign of saturation. These results indicate that V1 neurons have a wide dynamic range and respond robustly to contrast above 100%.

**Figure 5.**
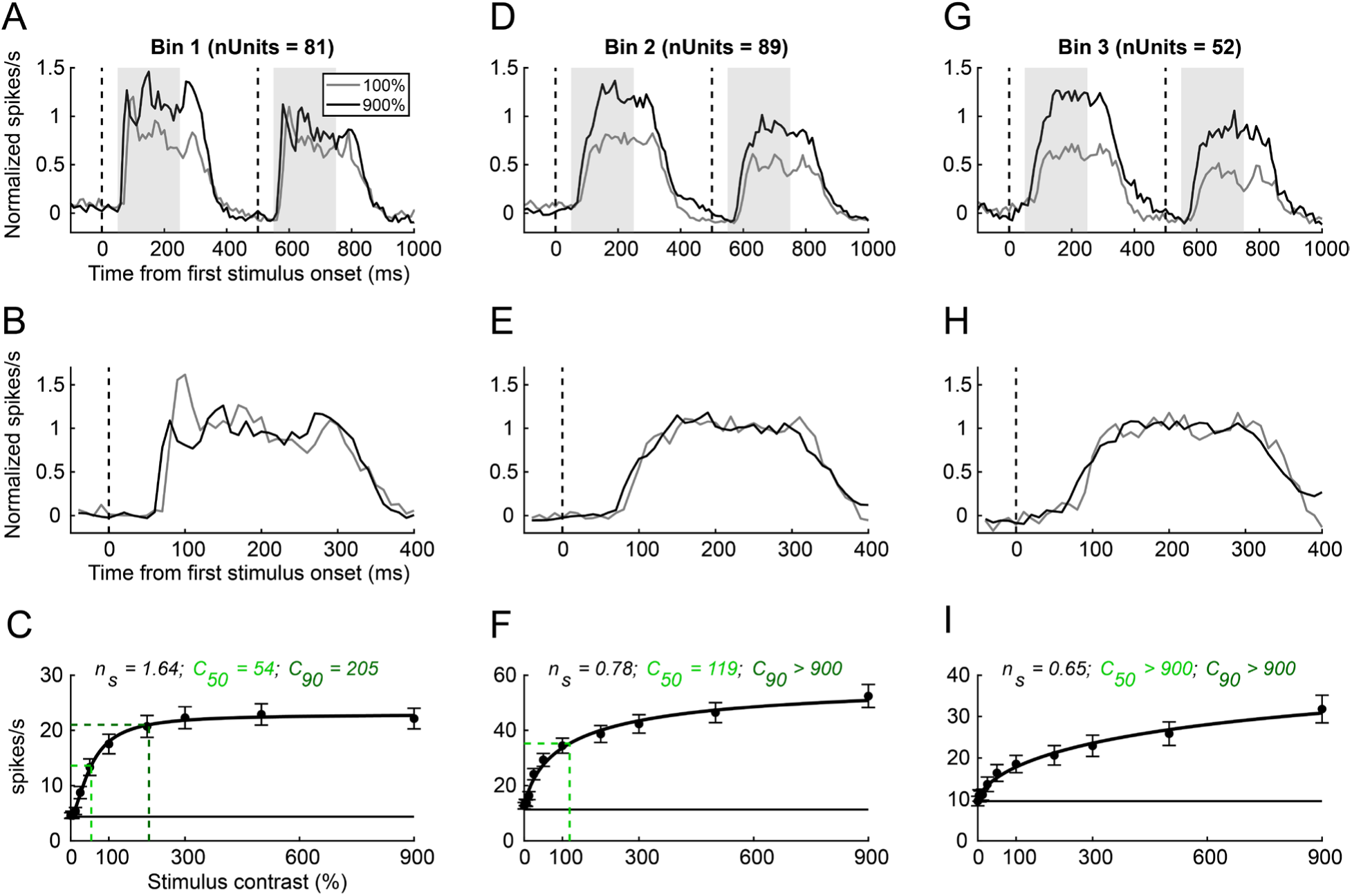
Average PSTH and CRF across subpopulations of single– and multi-units. *A,* Normalized average PSTHs across single– and multi-units in Bin 1 shown in Figure 4A. Same convention as Figure 3A. Spike count in the preceding 10 ms window is scaled to spike rate for every 10 ms and averaged across the units. The average PSTHs are normalized by the mean response between 100–300 ms after each stimulus onset at 900% contrast. ***B***, Same as ***A***, displayed over the first response cycle (0–400 ms) and normalized independently for each contrast level. ***C,*** Average CRF across the units in Bin 1. Error bars indicate SE across units. Same convention as Figure 3C. ***D-F*,** Same as ***A-C*** for Bin 2. ***G-I*,** Same as ***A-C*** for Bin 3.

### Population response in V1 and comparison with average unit spiking response

Our next goal was to determine whether the wide dynamic range of contrast-encoding in V1 observed with extracellular electrophysiology extends to measures of population responses using widefield calcium imaging (GCaMP). In a previous study, we found that GCaMP signals in V1 are approximately linearly related to the local spiking activities pooled across individual neurons (Seidemann et al., 2016). We therefore expected the dynamic range of the GCaMP signal to be similar to that of the pooled electrophysiological signals. To test this, we performed widefield GCaMP imaging in V1 in the cortical area encompassing the GCaMP expression site while the two animals viewed Gaussian stimuli across the same wide range of visual contrasts as in the electrophysiological recordings (see Methods). The stimuli were presented at the corresponding retinotopic coordinates of the GCaMP expression site.

To examine the dynamics and magnitude of the GCaMP response, we pooled the GCaMP signals over an approximately 3 × 2 mm^2^ region at the center of the cortical area activated by the Gaussian stimuli (see Methods). **Figure 6A and C** show the GCaMP responses to 100% and 900% contrast Gaussians in two example experiments. Consistent with our electrophysiological results, the GCaMP response to 900% contrast is much higher than the response to 100% contrast Gaussian.

**Figure 6.**
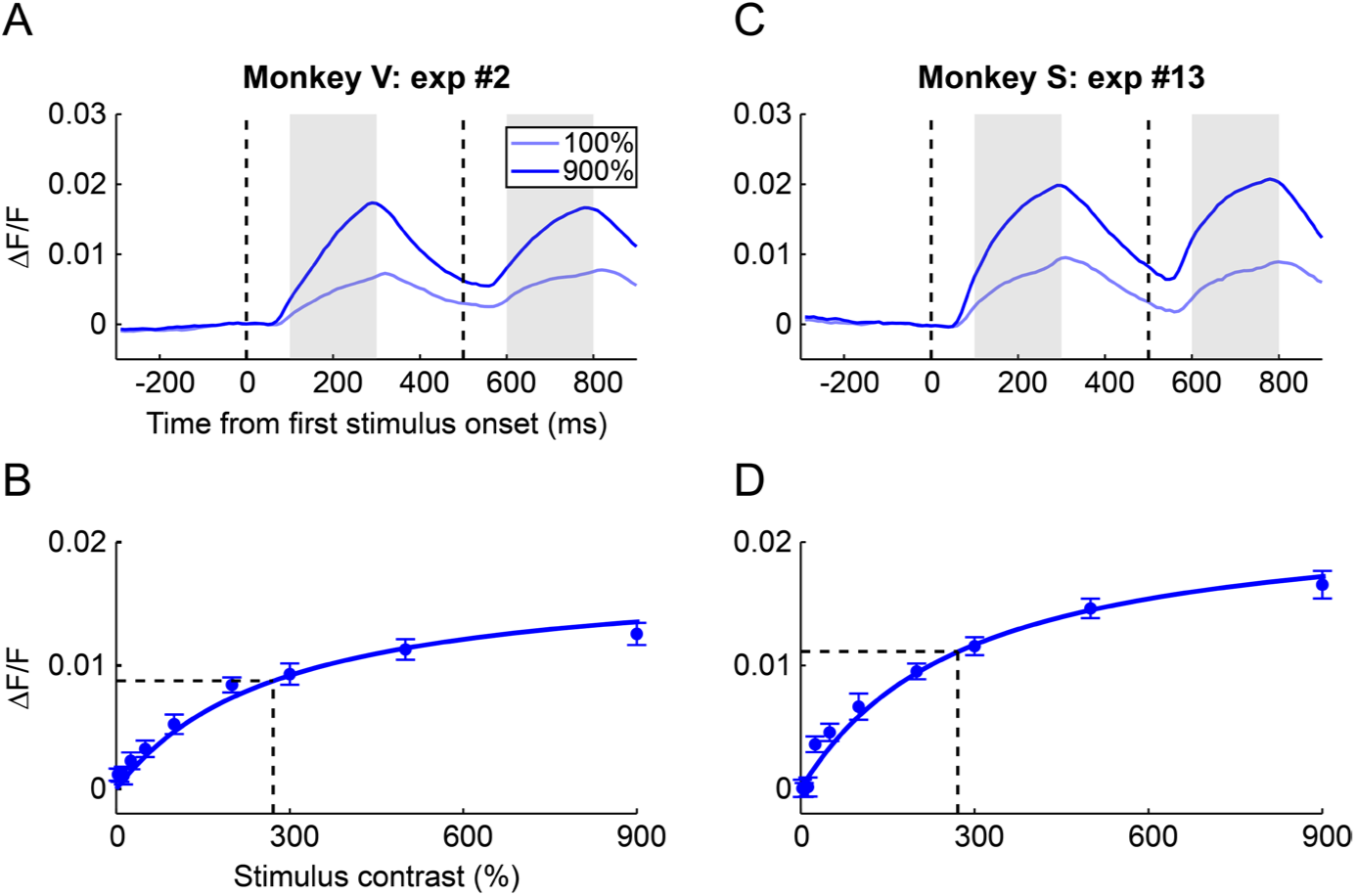
Example calcium population responses in V1 across a wide range of visual contrasts. *A,* Trial-averaged time courses of GCaMP responses to 100% and 900% contrast stimuli from an example experiment in Money V, aligned to the first stimulus onset. Same convention as Figure 3. The response is averaged over 100–300 ms after each stimulus onset to generate the CRF shown in ***B***. ***B,*** Trial-averaged CRF from the example experiment shown in ***A***, fitted by a Naka-Rushton curve (*n_s_*= 1.02, *C*_50_ = 271, *C*_90_ > 900). Error bars indicate SE across trials (n = 20). Dashed lines indicate the responses at *C*_50_. ***C-D,*** Same as ***A-B*** for an example experiment in Monkey S. The fitted curve in ***D*** shares the same parameters as the curve in ***B*** (see Methods).

**Figure 7.**
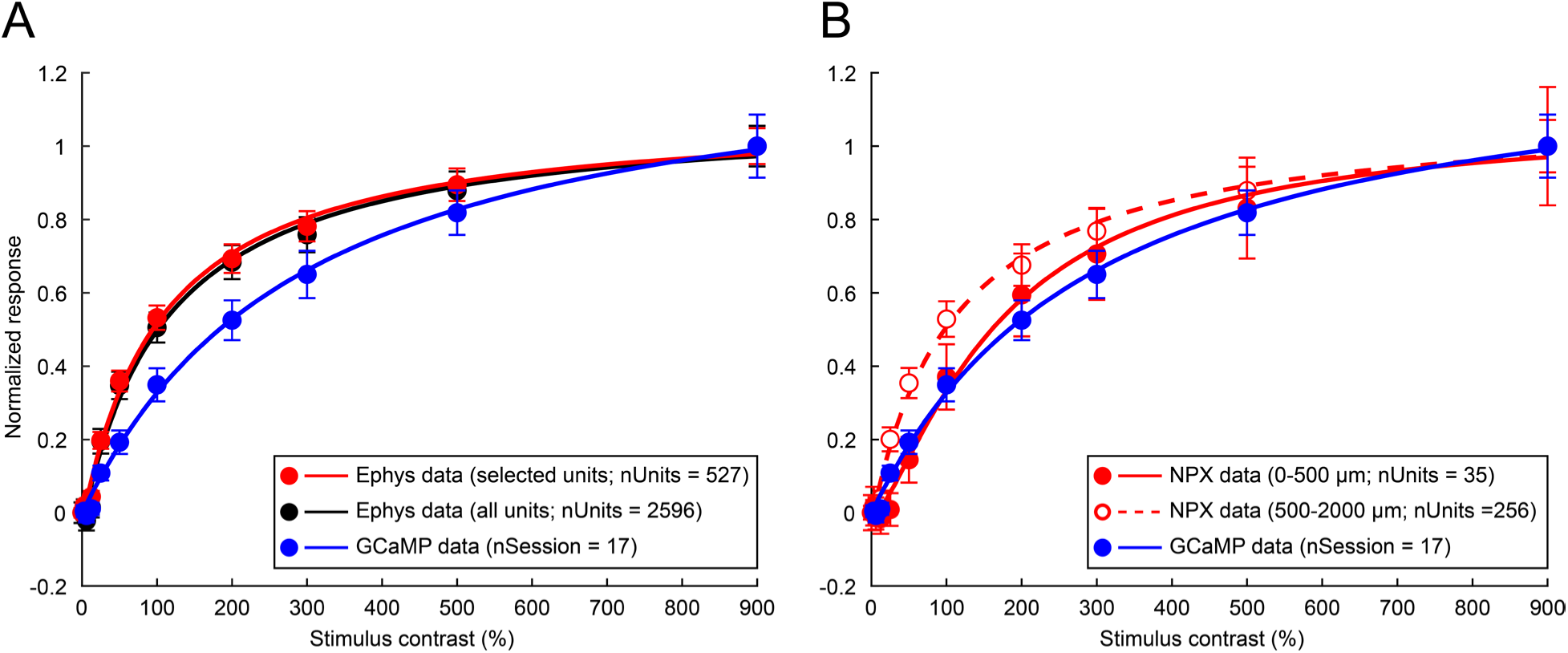
Comparison of electrophysiology data and imaging data. *A,* Spike rate or ΔF/F data is averaged across units or experiments and normalized by the mean response at 900% contrast. Curves are fitted to the mean responses across all the identified units (black; *n_s_*= 1.00, *C*_50_ = 112.64), the selected units (red; *n_s_* = 1.01, *C*_50_ = 106.22), and the imaging experiments (blue; *k_s_* = 1.02, *C*_50_ = 290.60). Error bars indicate SE across units or imaging experiments. ***B,*** Among the selected units, those recorded with Neuropixels probes are divided into two groups according to the cortical depths at which they were identified (solid red; *n_s_*= 1.37, *C*_50_ = 172.45; dashed red; *n_s_* = 0.98, *C*_50_ = 110.11).

To quantify the CRF of these responses, we computed the average amplitude of the GCaMP responses as a function of contrast (**Figure 6B and D**). The population GCaMP response continued to increase robustly beyond 100% contrast. Because we did not observe large variations in the shape of the CRF across experiments and monkeys, we fit all the GCaMP CRFs with a Nakka-Rushton function with shared *n_s_* and *C*_50_ while allowing *R_max_* to vary (to account for the varying imaging quality between experiments). The GCaMP CRFs had a small *n_s_* and a large *C*_50_, similar to one of our multi-unit examples (**Figure 3F**).

To compare the GCaMP and electrophysiology CRFs more directly, **Figure 7A** shows the average normalized CRF across all the selected units and the average CRF across all the calcium imaging experiments. Here, the Naka-Rushton function was fitted to the mean spiking response across all the single-and multi-units or the mean GCaMP responses across all the experiments. On average, the spiking response of the single– and multi-units reached only approximately 50% of its saturation at 100% contrast (when the data was normalized by the mean response at 900% contrast) and continued to increase robustly up to near 500% contrast where it reached approximately 90% of its saturation (**Figure 7A, red**). To examine whether our selection criteria biased our observed CRF, we repeated the analysis without excluding any units. The average CRF of all units (**Figure 7A, black**) was indistinguishable from that of the selected subset, indicating that our results are robust to the selection criteria of the electrophysiology data set.

The increase in the visual response above 100% contrast was even more pronounced in the GCaMP signals (**Figure 7A, blue**). The average GCaMP response reached only approximately 30% of its saturation at 100% contrast, creating a mismatch with the average unit spiking response. Since widefield GCaMP signals are strongly dominated by neurons in superficial layers in the upper 500 µm of the cortex (Yizhar et al., 2011), we divided the selected single– and multi-units into two groups according to their cortical depths and compared their average CRFs to the GCaMP data (**Figure 7B**). The analysis was limited to the Neuropixels recordings where we had a more accurate estimate of the depth of each recorded unit. The average CRF of units recorded at depths of 0–500 µm had a higher *C*_50_ and aligned more closely with the GCaMP data than the average CRF of units recorded at 500–2000 µm. These results are generally consistent with our prior findings that widefield GCaMP signals are approximately linearly related to the local spiking activities pooled across individual neurons in superficial cortex. Overall, our electrophysiological and GCaMP results show that V1 neural populations have a wide dynamic range of contrast encoding.

## Discussion

Previous measurements of contrast response functions (CRFs) in primary visual cortex (V1) have been constrained to Weber contrasts less than or equal to 100%. We measured local Weber contrasts in natural scenes and found that a substantial percentage of patches exceed 100% contrast (**Figure 1**). Given the prevalence of high contrasts in natural scenes, it is plausible that V1 encodes a much larger range of Weber contrasts than previously thought. To test this hypothesis, we measured single-unit, multi-unit, and GCaMP responses to small Gaussian targets, for Weber contrasts ranging from 0% to 900%.

Our sample of single– and multi-units in V1 exhibited a range of CRF shapes, which were quantified by the two shape parameters of the Naka-Rushton function: the semi-saturation constant *C*_50_, and the exponent *n_s_*. While some units reached 90% of their maximum response at a contrast (*C*_90_) below 100%, the vast majority of units did not saturate until the contrast substantially exceeded 100%, supporting the hypothesis that many V1 neurons usefully encode contrasts well above 100% Weber contrast. After excluding those with *C*_90_ > 900, the mean *C*_90_ remained above 100% for both single– and multi-units. Among the remaining units, those with *C*_90_ ≤ 100 tended to have a higher exponent (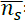 = 2.41), while those with *C*_90_ > 100 tended to have a lower exponent (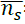 = 1.66).

These findings show that V1 contains neural subpopulations with a broad contrast dynamic range extending to contrast well above 100% Weber contrast. The observed systematic relationship between the power exponent, *n_s_*, and *C*_50_ shows that neurons that continue to respond robustly to Weber contrasts above 100% (i.e., high *C*_50_ neurons) tend to have exponents *n_s_* closer to 1, which means that their fitted Naka-Rushton curves to are relatively linear in the low contrast range (i.e., below 100%), as can be seen in the CRFs of the example units and the average CRFs within the neural subpopulations (**Figure 3D and F, and Figure 5F and I**). However, it is also important to note that the typical accelerating rise often observed in the low contrast range might be missed when fitting such a wide contrast range (i.e., Naka-Rushton function may not be a sufficiently accurate description over the full dynamic range).

We estimated population activity in V1 solely from the spiking activity of neurons, by averaging the CRFs across our selected sample of single– and multi-units and compared it to the average GCaMP response, which is a measurable correlate of population activity. Consistent with the fact that most single– and multi-units had a *C*_90_ greater than 100% Weber contrast, the average unit response continued increasing above 100%. Also, the exponent of the average unit response is 1.01, and hence the contrast response is approximately linear over the lower range of contrasts. Similarly, the average GCaMP response continued to increase robustly across the high contrast range. We also note that previous work suggests Gabor and Gaussian stimuli of equivalent maximal luminance evoke comparable V1 population responses (Chen et al., 2012). Thus, our results strongly suggest that the dynamic range of V1 population responses to localized stimuli is substantially wider than previously assumed.

However, the increase in the response across the high contrasts was even more pronounced in the GCaMP signals, creating a significant discrepancy between the two average CRFs (**Figure 7A**). This result differs from our previous study, which showed that widefield GCaMP signals are approximately linearly related to the locally pooled spiking activity (Seidemann et al., 2016). Several factors may contribute to this difference, with the first being the variation in the neural population recorded by the two methods. While the widefield imaging predominantly captures neural activity in superficial layers of the cortex (up to the depth of ∼500 µm) due to the limited penetration depth of the blue excitation light (Yizhar et al., 2011), laminar probes used in the present study record activity from not only superficial, but also deeper layers. Thus, the overall contrast response profile of the neural population could differ between the two recording methods, resulting in the mismatch. Consistent with this, our analysis of the Neuropixels recordings shows that the average CRF of units recorded at cortical depths of 0–500 µm has a higher *C*_50_ and aligns more closely with the GCaMP signals than the average CRF of units recorded beyond 500 µm. These results suggest that the neuronal pool in superficial layers (L1 and L2/3) is dominated by neurons with a high *C*_50_, while the pool in deeper layers contains a higher proportion of neurons with a low *C*_50_. In addition, our sampling of single– and multi-units may simply be biased towards a subpopulation of neurons that start to saturate at lower contrasts (i.e., low *C*_50_), regardless of their cortical depth. In addition, while the calcium signal was captured solely from the excitatory neurons (targeted with the CaMKIIa promoter; see Methods), the electrophysiology data was collected from both excitatory and inhibitory neurons.

The discrepancy could also represent the nonlinear relationship between spiking of single neurons and GCaMP fluorescence (Akerboom et al., 2012; Chen et al., 2013). GCaMP signals provide an indirect measure of spiking activity by reflecting the changes in the intracellular calcium concentrations. However, the transformation from spiking activity to calcium levels is inherently nonlinear due to the subthreshold activity and calcium dynamics (Scheuss et al., 2006). The transformation from calcium to GCaMP fluorescence is also nonlinear due to the fluorescence kinetics and the calcium binding affinity of the GCaMP protein (Zhang et al., 2023; Zhang and Looger, 2023). These calcium nonlinearities in GCaMP signals may have persisted despite the pooling of heterogeneous neurons.

In previous work (Chen et al., 2022), we found that direct optogenetic stimulation in V1 can produce population responses substantially greater than those produced by 100% contrast Gaussian stimuli. One hypothesis for this finding is that the optogenetic stimulation was pushing the neural population response beyond its normal operating range. However, the current findings suggest that this may not be true. Further testing with higher contrast visual stimuli will be required to compare the population dynamic ranges of visual and optogenetic stimulation.

Our current study comes with several limitations. First, the quality of preprocessing of our electrophysiology data and isolation of visual responses are inherently constrained by the algorithm of Kilosort3 and potential human errors in manual spike-sorting. In our main analysis, we excluded the majority of our single– and multi-unit samples because of their noisy and unreliable visual responses. Nevertheless, we demonstrated that the shape of the contrast response function of the pooled spiking activity is largely unaffected by including the entire sample of units (**Figure 7A**). Second, while the size of the visual stimulus was kept small, it was not optimally tuned to the receptive field size of the recorded neurons, leaving open the possibility of surround suppression (Sceniak et al., 1999). Moreover, the stimulus position was not always optimized, which may have caused the stimulus to be offset from the receptive field center of the recorded neurons.

In conclusion, the dynamic range of single neurons and neural populations in V1 are substantially greater than previously assumed. The wider dynamic range also aligns more closely with the statistics of contrast in natural scenes, suggesting that there is evolutionary value in encoding the high contrasts found in the natural environment.

### Future directions

Here, we reveal that the dynamic range of contrast-encoding in V1 population responses to localized stimuli extends beyond the commonly presumed range. However, the present finding may not hold at the single-cell level. Past studies often recorded the spiking responses of individual neurons using Gabor or grating stimuli optimized for their preferred orientation and spatial frequency, whereas the Gaussian stimulus used here is suboptimal for driving the individual neurons. An important future direction is to compare single-neuron responses to Gabor and Gaussian stimuli. For instance, one can investigate whether the response to Gabor stimulus starts to saturate below 100% Michelson contrast in neurons with a wide dynamic range and whether a Gaussian of extremely high Weber contrast (e.g., 900%) can drive these neurons to fire more strongly than an optimally tuned Gabor of 100% Michelson contrast. Additionally, a high-contrast stimulus that contains both the orientation and spatial frequency information can be generated by summing a Gaussian and a Gabor of the same amplitude. This composite stimulus, referred to as “increment-Gabor” by Kortum and Geisler (Kortum and Geisler, 1995), provides a valuable tool for investigating the contributions of different visual features to the activity of individual neurons and the population response in V1.

The present study has only focused on the neural representation of contrast encoding in V1. An additional future step should be taken to measure discrimination of Weber contrasts in the animals and compare their detection thresholds with those computed from the population responses, to determine whether the animals’ behavior aligns with the measured population responses.

## Methods

### 1. Natural scene images and center-surround contrasts

We computed the distribution of the center-surround contrast, *C_cs_* in five 14-bit natural scene images (50 pixels per degree) by converting them into grayscale intensity images (16,384 gray levels) and convolving them with kernels consisting of non-overlapping concentric center and surround subregions at each image pixel location. To compute *C_cs_* at a given image pixel location (*x*, *y*), a kernel was placed on the image such that its central pixel aligned with the image pixel. The kernels are weight vectors in which each of the *N* pixels in the center and surround regions is assigned a uniform weight of 1/*N*, computed independently for each region. Convolution of the image with the kernel yields the mean luminance of the image pixels falling within the center (*L_c_*_(*x*, *y*)_) and surround regions (*L_s_*_(*x*, *y*)_) that are required to calculate (*C_cs_*_(*x*, *y*)_).

### 2. Recording chamber and viral injection

We implanted two male rhesus macaques (*Macaca mulatta;* Monkey V and Monkey S) with a custom metal recording chamber over dorsal V1 in the right hemisphere to gain access to a cortical area representing the lower-left visual field. We injected 4–5 μl of a GCaMP6f viral vector into the cortex per site at depths up to 2000 µm (Monkey V with AAV1-CaMKIIa-GCaMP6f, and Monkey S with AAV1-CaMKIIa-GCaMP6f or AAV1-CaMKIIa-NES-GCaMP6f) based on the methods from our previous study (Seidemann et al., 2016).

### 3. Fixation task and stimulus contrasts

We trained the monkeys to perform a fixation task where they were required to maintain fixation at the center of the screen in the presence of a small flashing Gaussian stimulus (0.33° FWHM). The task and visual stimulus were presented on a Sony CRT monitor (1024 × 768 @100 Hz) positioned 108 cm from the eyes, yielding a stimulus resolution of 50 pixels per degree. Eye position was monitored with an EyeLink 1000 Plus system. Background luminance of the monitor was lowered to 10% of its maximum (i.e., 10 cd/m^2^) to create a uniform dark grey background.

For Monkey S, whose retinotopy was estimated by voltage-sensitive dye imaging (Yang et al., 2007), the stimulus was placed at the corresponding retinotopic coordinates of the GCaMP expression site in the imaging area or the probe penetration site. For Monkey V, for which retinotopy was not available, we used the stimulus with high Weber contrast (900%) to estimate the receptive field center of the neurons in the recording site.

During experiments, the animals were seated in a custom chair with their head immobilized via a head-post implant. Each trial began with fixation on a small white 0.1° × 0.1° square at the center of the screen. After an initial fixation period of ∼1300 ms (pre-stimulus fixation duration was varied across trials), the stimulus was presented at 2 Hz (200 ms ON and 300 ms OFF for 2 cycles). The animals were rewarded for continuing to maintain fixation for 1200 ms after the first stimulus onset. Behavioral data were collected with custom MATLAB-based software.

### 4. Electrophysiology

We recorded extracellular spiking activities of V1 neurons using multi-channel probes (Plexon S-probes and NHP Neuropixels probe 1.0). To prevent large drifts of the probe in the brain during the recording, we applied a small pressure on the cortex with a custom plastic insert and left the probe in the brain for 60-90 minutes before data collection. The probe was always inserted approximately normal to the brain surface. We advanced the probe into the brain until all the electrode patches were placed in the brain. Real-time spiking data were collected using Trellis and SpikeGLX software for the Plexon and Neuropixels sessions, respectively. The visual stimulus was placed at 1.4–2.7° eccentricity from the fixation point in the lower left quadrant of the visual field.

### 5. Analysis of electrophysiology data

The electrophysiological data were first spike-sorted automatically into individual single– or multi-unit clusters by Kilosort3. By utilizing the statistical analysis (principal component analysis and autocorrelation analysis of spikes, and distribution of spike amplitudes) generated by Phy, we manually approved or disapproved the automatic sorting of spike clusters. Similar single-unit spike clusters were merged, and dissimilar spikes were either split or excluded.

For each single– or multi-unit in a given trial, we computed the spike counts over 50–250 ms after the onset of each stimulus cycle, scaled them to spike rate (i.e., number of spikes per second). The spike rates were averaged across two stimulus cycles for each trial and averaged across trials for each condition.

The spiking response of each identified single– or multi-unit as a function of stimulus contrast (*R_spike_*(*c*)) was fitted independently by the following Naka-Rushton function:

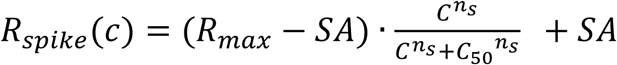

where *C*_50_ is the semi-saturation constant, *n_s_* is the power exponent, *SA* is the spontaneous firing rate and *R_max_* is the maximum firing rate. The parameters were fitted independently for each unit by maximum-likelihood estimation (MLE) method. The power exponent, *n_s_* was fitted be between 0 and 3 and the semi-saturation constant, *C*_50_ was fitted between 0 and 900. Only the units with *R_max_* ≥ 5, *R_max_*/*SA* ≥ 3, and *R*^2^ > 0.8 were included for analysis. *R*^2^ is the coefficient of determination. The near-saturation contrast (*C*_90_) of each unit was extrapolated from the fitted curve. To compare with the GCaMP data, we averaged the spiking responses across all the selected units and fitted the mean unit spiking response with the same function.

To describe the relationship between *C*_50_ and *n_s_* across all the selected units, we fitted the following exponential decay function where *k*_1_ is the scaling constant, *m* is the decay rate constant, and *k*_2_ is the additive constant which describes the lower asymptote that the function approaches as *n_s_* increases towards infinity.

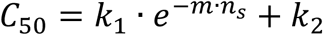

### 6. Calcium imaging

We performed widefield calcium imaging using the procedures described previously (Chen et al., 2022; Seidemann et al., 2016). The imaging system comprised of a PCO Edge CLHS sCMOS camera, a 50 mm fixed-focus objective lens (facing cortex), a dichroic mirror (long-pass at 505 nm) and an 85 mm fixed-focus objective lens. Calcium signals were captured at a frame rate of 100 Hz from an 8.2 × 8.2 mm^2^ cortical area approximately centered on the GCaMP expression site. Images were binned to 512 × 512 pixels. Excitation light (X-Cite 120 LED) and GCaMP emission were bandpass-filtered at 480 nm and 520 nm, respectively. Data were collected with custom MATLAB-based software. Data acquisition was time-locked to the heartbeat using EKG (HP Patient Monitor) to enable the removal of the heartbeat artifact in offline data analysis. The visual stimulus was placed in the lower-left visual field at 1.3° eccentricity from the fixation point for Monkey V and 2.3 or 2.7° eccentricity for Monkey S.

### 7. Analysis of imaging data

The imaging data were down-sampled and normalized to compute the physiological signal expressed as Δ*F*/*F*. For both monkeys, the average florescence over frames 0-300 ms prior to stimulus onset was used as the baseline, *F*_0_.

To extract visual-evoked response, the signal was first averaged temporally over frames 100-300 ms after each stimulus cycle onset and anchored by the frames 0-100 ms prior to first stimulus cycle onset for each trial. The signals were then averaged spatially across the selected ROI (3.1 × 2.4 mm^2^ cortical area for Monkey V, and 2.3 × 1.5 mm^2^ or 2.8 × 1.9 mm^2^ cortical area for Monkey S) and across trials for each condition. Finally, the mean response in blank condition was subtracted from the mean response of each experimental condition to remove the heartbeat artifact.

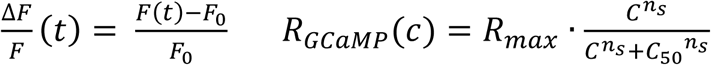

For each experiment, the GCaMP response as a function of stimulus contrast (*R_GCaMP_*(*c*)) was fitted by the same function used to fit the electrophysiological data; however, the spontaneous firing rate, *SA* was removed from the equation due to the blank subtraction. The fitted maximum response, *R_max_* allowed the function to account for the variability in the amplitude of signal and imaging quality across recording blocks or sessions due to various factors including, but not limited to, the difference in the GCaMP expression across the animals, layer regrowth over the cortex, and discharge that affects the clarity of the cortical chamber.

The parameters were fitted by MLE method. To fit the GCaMP response in each experiment, the power exponent, *n_s_* and the semi-saturation constant, *C*_50_ were forced to be shared across blocks, while *R_max_* was fitted independently for each block. To compare with the electrophysiology data, we averaged the GCaMP responses across experiments and fitted the mean GCaMP response with the same function.

